# Cross-modal Suppression of Auditory Association Cortex by Visual Speech as a Mechanism for Audiovisual Speech Perception

**DOI:** 10.1101/626259

**Authors:** Patrick J. Karas, John F. Magnotti, Brian A. Metzger, Lin L. Zhu, Kristen B. Smith, Daniel Yoshor, Michael S. Beauchamp

**Affiliations:** Department of Neurosurgery, Baylor College of Medicine

## Abstract

Vision provides a perceptual head start for speech perception because most speech is “mouth-leading”: visual information from the talker’s mouth is available before auditory information from the voice. However, some speech is “voice-leading” (auditory before visual). Consistent with a model in which vision modulates subsequent auditory processing, there was a larger perceptual benefit of visual speech for mouth-leading *vs.* voice-leading words (28% *vs.* 4%). The neural substrates of this difference were examined by recording broadband high-frequency activity from electrodes implanted over auditory association cortex in the posterior superior temporal gyrus (pSTG) of epileptic patients. Responses were smaller for audiovisual *vs.* auditory-only mouth-leading words (34% difference) while there was little difference (5%) for voice-leading words. Evidence for cross-modal suppression of auditory cortex complements our previous work showing enhancement of visual cortex (Ozker et al., 2018b) and confirms that multisensory interactions are a powerful modulator of activity throughout the speech perception network.

**Impact Statement:** Human perception and brain responses differ between words in which mouth movements are visible before the voice is heard and words for which the reverse is true.

## Introduction

Pairing noisy auditory speech with a video of the talker dramatically improves perception (Bernstein et al., 2004; Grant and Seitz, 2000; Munhall et al., 2004; Ross et al., 2007; Sumby and Pollack, 1954). As shown in Figure 1A, the preparatory mouth movements of visual speech begin before auditory vocalization (Figure 1A) providing both an alert about impending auditory speech and information about the expected speech content. Since any particular visual mouth movement is compatible with only a few auditory phonemes, perceptual accuracy can be improved by suppressing cortical representations of incompatible phonemes and enhancing representations of compatible phonemes in the interval after visual speech begins but before auditory speech arrives (Cappelletta and Harte, 2012; Jeffers and Barley, 1971; Neti et al., 2000).

**Figure 1.**
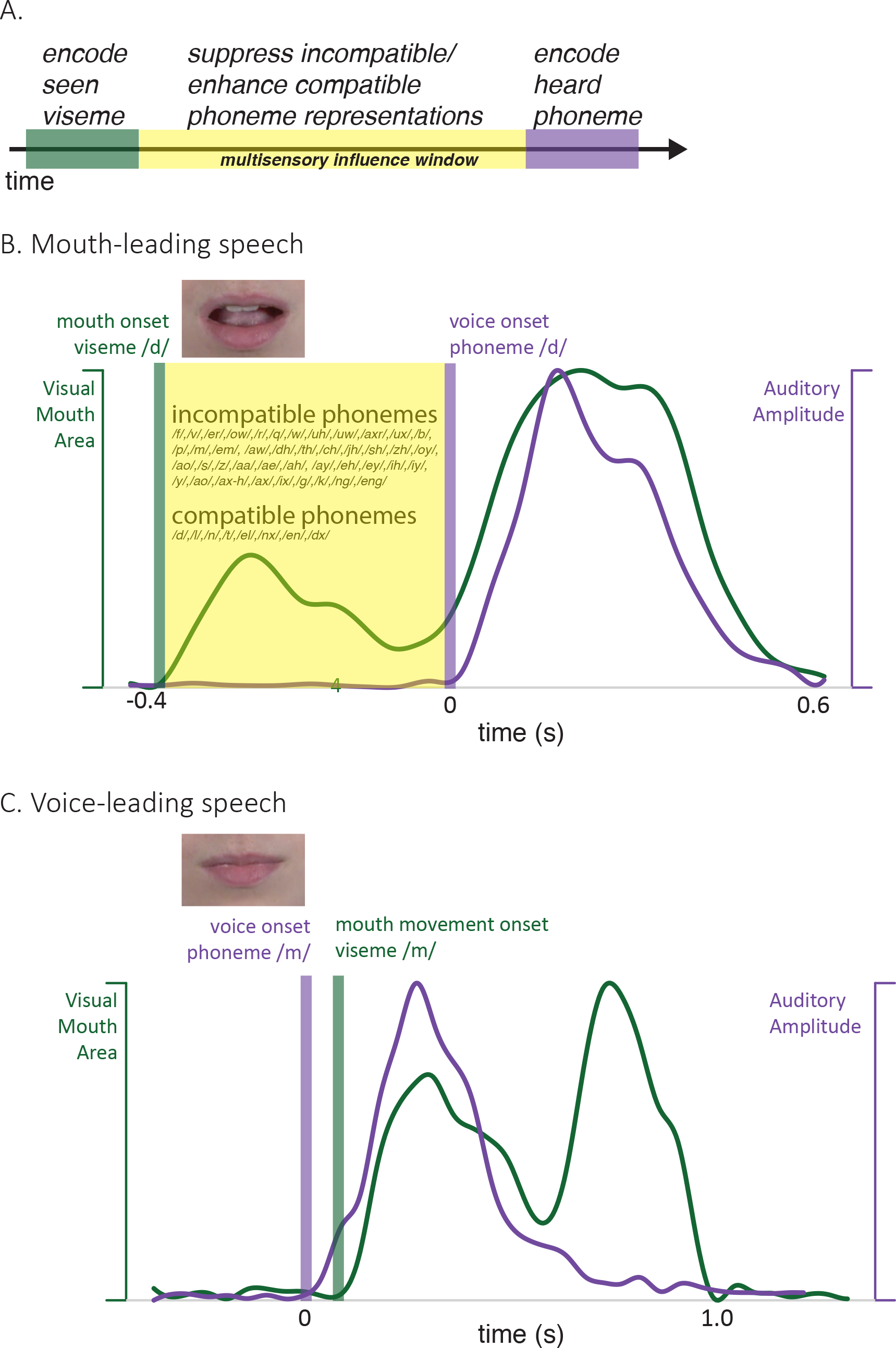
Time course of audiovisual speech. A. Conceptual model of audiovisual speech showing the onset of visual speech features (visemes, green) prior to the onset of auditory speech features (phonemes, purple) creating a multisensory influence window during which compatible phoneme representations can be enhanced and incompatible phoneme representations suppressed (yellow), improving perception.B. Measurements of auditory and visual speech feature asynchrony for the word “drive.” Audiovisual speech is composed of visual mouth movements (green line showing visual mouth area) and auditory speech sounds (purple line showing auditory sound pressure level). For the word “drive,” lip and mouth movements (visual speech onset, green bar) occur prior to vocalization (auditory speech onset, purple bar). Time zero is the auditory speech onset. This speech is termed “mouth-leading” as visual mouth movements begin before auditory speech. C. For the word “known,” mouth movements begin after auditory vocalization (green bar comes after purple bar). This speech is termed “voice-leading” as vocalization begins before visible mouth movements.

Perceptual studies on the benefits of visual speech are experimentally straightforward: accuracy is compared between conditions without visual speech and with visual speech added. This straightforward approach is problematic for neural studies because viewing a moving face activates a host of brain regions and processes, many of which may not contribute to speech perception. We tackle this difficulty with an approach that leverages natural variability in the temporal relationship between modalities (Chandrasekaran et al., 2009; Schwartz and Savariaux, 2014). Most audiovisual speech is “mouth-leading”: the visual information provided by the talker’s mouth is available before auditory information from the talker’s voice. For an audiovisual recording of the word “drive” (Figure 1B) the visual onset of the open mouth required to enunciate the initial “d” of the word preceded auditory vocalization by 400 ms, allowing the observer to rule out the ~80% of phonemes incompatible with this mouth shape (Cappelletta and Harte, 2012). However, some audiovisual speech is “voice-leading”: auditory speech precedes visual speech. For an audiovisual recording of the word “known”, visual changes did not begin until almost 100 ms *after* auditory voice onset (Figure 1C).

If the perceptual benefit of visual speech results from the head start that it provides for auditory processing, this benefit should be reduced or eliminated for voice-leading speech, in which the visual modality does not provide speech information until after the auditory modality. Any perceptual benefit or lack thereof is expected to be mirrored in measurements of neural activity. Neural responses specific to mouth-leading words can then be attributed to multisensory integration rather than non-specific responses to the presence of a moving face, which is present in both mouth-leading and voice-leading stimuli.

Studies of patients with cortical lesions (Hickok et al., 2018; Stasenko et al., 2015) and fMRI, MEG, EEG and electrocorticographic (intracranial EEG) studies have shed light on the neural mechanisms underlying audiovisual speech perception, implicating a network of brain areas in occipital, temporal, frontal and parietal cortex (Crosse et al., 2016; Hickok and Poeppel, 2015; Okada et al., 2013; Salmelin, 2007; Shahin et al., 2017; Sohoglu and Davis, 2016; van Wassenhove et al., 2005). Within this network, posterior superior temporal gyrus and sulcus (pSTG) is responsive to both unisensory auditory speech (Belin et al., 2000; Formisano et al., 2008; Mesgarani et al., 2014) and unisensory visual speech (Bernstein et al., 2011; Bernstein and Liebenthal, 2014), with subregions responsive to both auditory and visual speech (Beauchamp et al., 2004; Ozker et al., 2017; Ozker et al., 2018a; Rennig et al., 2018; Zhu and Beauchamp, 2017). We examined the neural differences between the processing of mouth-leading and voice-leading speech in pSTG using intracranial electrocorticography. This technique has the advantage of high spatial resolution (necessary to measure activity from focal areas within the pSTG) and high temporal resolution (necessary to capture the small auditory-visual asynchrony differences between visual-leading and mouth-leading words).

## Results

### Perceptual Results

We examined the perception of auditory-only and audiovisual words in 40 participants. Perception was very accurate (mean accuracy 95%) for clear words without added auditory noise. Adding noise reduced perceptual accuracy below ceiling, revealing differences between conditions (Figure 2A). For mouth-leading words, viewing the face of the talker increased the intelligibility of noisy auditory speech by 28%, from 32% for auditory-only words to 60% for audiovisual words. For voice-leading words, viewing the face of the talker provided only a 4% increase, from 77% to 81%.

**Figure 2.**
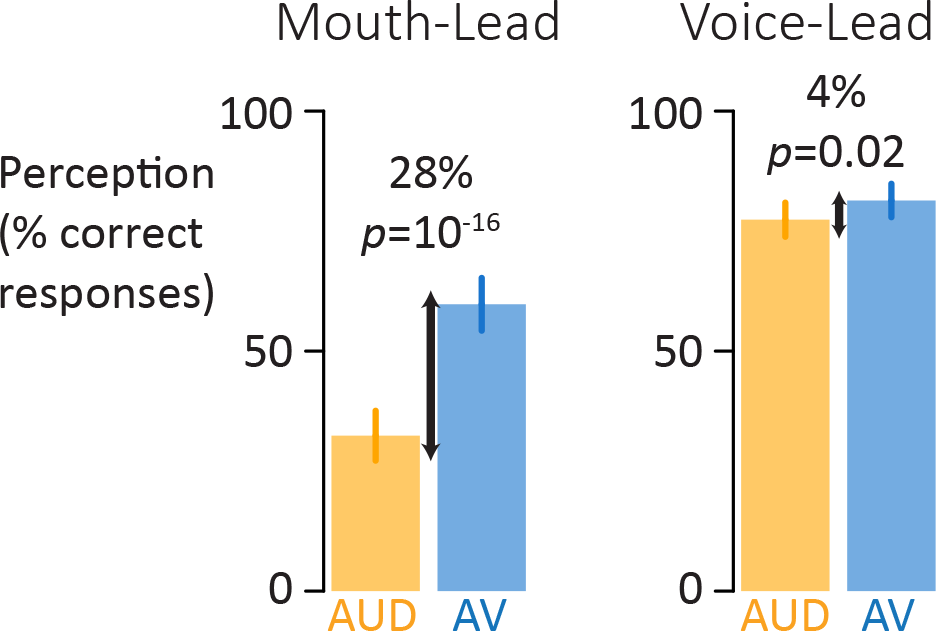
Perceptual performance on speech-in-noise recognition task. A. For mouth-leading words (left plot), the addition of visual speech increased comprehension of speech-in-noise words by 28% (black arrow), from 32% correct for auditory-only speech (orange) to 60% correct for audiovisual speech (blue). In contrast, for voice-leading words (right plot) the addition of visual speech increased accuracy by 4%, from 77% for auditory-only speech to 81% for audiovisual speech. Error bars represent standard error of the mean across subjects.

To evaluate these differences, we constructed a generalized linear mixed-effects model (GLMM) with fixed effects of word format (auditory-only *vs*. audiovisual) and word type (mouth-leading *vs.* voice-leading) and their interaction. The GLMM showed a significant interaction between format and word type (*p* = 10^−9^) driven by a greater accuracy improvement between the auditory and audiovisual formats for mouth-leading words (28% accuracy increase, odds-ratio 6.1, *p* < 10^−16^) than for voice-leading words (4% accuracy increase, odds-ratio 1.4, *p* = 0.02). In addition to the interaction, there were significant main effects of format (*p* < 10^−16^) and word type (*p* = 0.01) driven by higher accuracy for audiovisual words and for voice-leading words.

### Neural Results

The perceptual results supported the conceptual model prediction of a greater multisensory benefit for mouth-leading words than voice-leading speech. To study the neural correlates of this difference, in eight epileptic participants we recorded from electrodes implanted bilaterally over the posterior superior temporal gyrus (pSTG) that showed a significant response to auditory-only speech measured as the percent increase in the power of the high-frequency (75 to 150 Hz) electrical activity relative to baseline (*n* = 28 electrodes, locations and auditory-only response magnitude shown in Figure 3A). In contrast to the perceptual studies, where both clear and noisy speech was presented, in the neural experiments only clear speech was presented.

**Figure 3.**
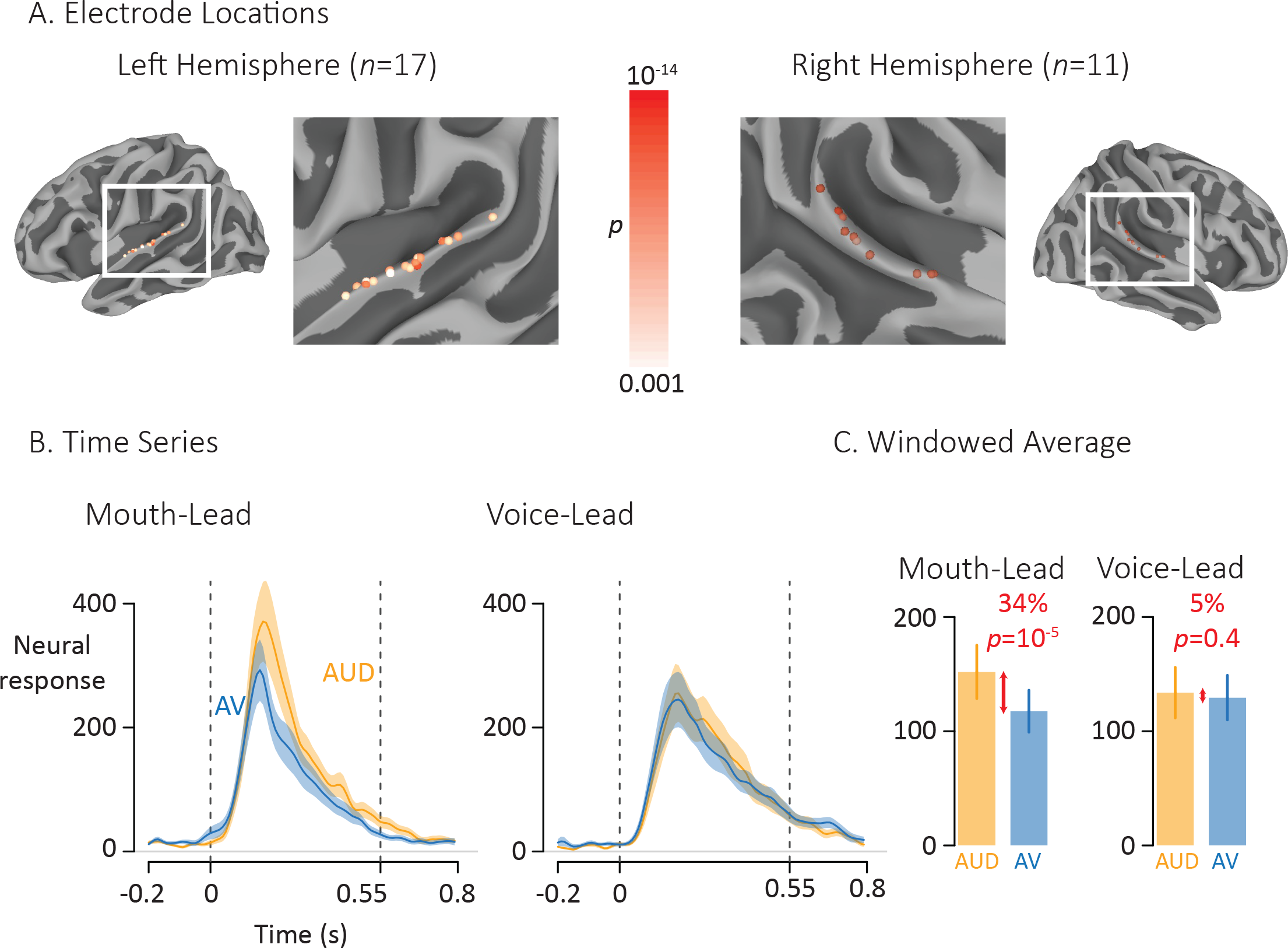
Average broadband high-frequency activity by experimental condition. A. The location of 17 left-hemisphere (left panel) and 11 right-hemisphere (right panel) electrodes that met both an anatomical criterion (located over the posterior superior temporal gyrus) and a functional criterion (significant response to auditory-only speech). The color of each electrode shows the significance (corrected for multiple comparisons using *p* < 0.001 Bonferroni-corrected) of each electrode’s response to the auditory-only condition during the period from auditory speech onset to offset. B. For mouth-leading words (left panel), the neural response to auditory-only words (AUD; orange line) was greater than the response to audiovisual words (AV; blue line). For voice-leading words (right panel), the responses were similar. Shaded regions show standard error of the mean across electrodes (*n* = 28) and dashed lines show auditory speech onset (0 s) and offset (0.55 s). C. To quantify the difference between word types, the neural response was averaged within the window defined by auditory speech onset and offset. For mouth-leading words (left panel), the auditory-only format evoked a significantly greater response than the audiovisual format (34% difference, *p* = 10^−5^). For voice-leading words, there was little difference (5%, *p* = 0.41), resulting in a significant interaction between word format and word type (34% *vs.* 5%, *p* = 0.001). Error bars show standard error of the mean across electrodes (*n* = 28).

As shown in Figure 3B, the neural response to the single word stimuli in the pSTG began shortly after auditory speech onset at 0 ms, peaked at 180 ms, and returned to baseline after auditory speech offset at 550 ms. For mouth-leading words, the response to audiovisual words (relative to baseline) was smaller than the response to auditory-only words. For voice-leading words, the responses to audiovisual and auditory-only words were similar, confirming the model prediction of a greater multisensory effect for mouth-leading than voice-leading words.

To quantify this effect, we entered the mean response across the window from auditory speech onset to offset (0 ms to 550 ms) into a linear mixed-effects model (LME) with fixed effects of word format (auditory-only *vs.* audiovisual), word type (mouth-leading *vs.* voice-leading) and the interaction (Figure 3C). The interaction was significant (*p* = 0.001) driven by a smaller increase from baseline for audiovisual *vs.* auditory-only for mouth-leading words (34%, 118% *vs.* 152%, *p* = 10^−5^) than for voice-leading words, (5%, 129% *vs.* 134%, *p* = 0.4). There were significant main effects of word format (*p* = 10^−5^, driven by higher amplitude for auditory-only) and word type (*p* = 0.005, driven by higher amplitude for mouth-leading).

### Multisensory Influence of Visual Speech: Mouth-Leading Words

The conceptual model posits both that visual speech information is available and that it arrives early enough to exert multisensory influence. To determine if these assertions were true in the pSTG, we examined the responses to visual-only words. There was significant positive response to visual-only words, with a mean amplitude of 20% (0 to 200 ms after visual speech onset; significantly greater than baseline, *p* = 10^−7^, single sample *t*-test). The effect was consistent across electrodes, with 27 out of 28 electrodes showing a positive response to visual-only speech, demonstrating that information about visual speech reaches pSTG. The model also requires that the visual speech information arrives early enough in the pSTG to exert multisensory influence. As shown in Figure 4A, the pSTG response to visual speech features occurred ~100 ms earlier than the response to auditory speech features, sufficient time for multisensory interactions to occur. To further quantify this observation, we calculated the latency of the response (defined as the time of half-maximum response) to visual-only and auditory-only speech in individual electrodes. Responses were aligned to auditory onset (or to the time when auditory onset would have occurred for visual-only stimuli), with the response to visual-only speech occurring a mean of 123 ms earlier than the response to auditory-only speech (paired *t*-test, *p* = 10^−8^). For 26 out of 28 electrodes, the response to visual-only mouth-leading words occurred earlier than the response to auditory-only mouth-leading words.

**Figure 4.**
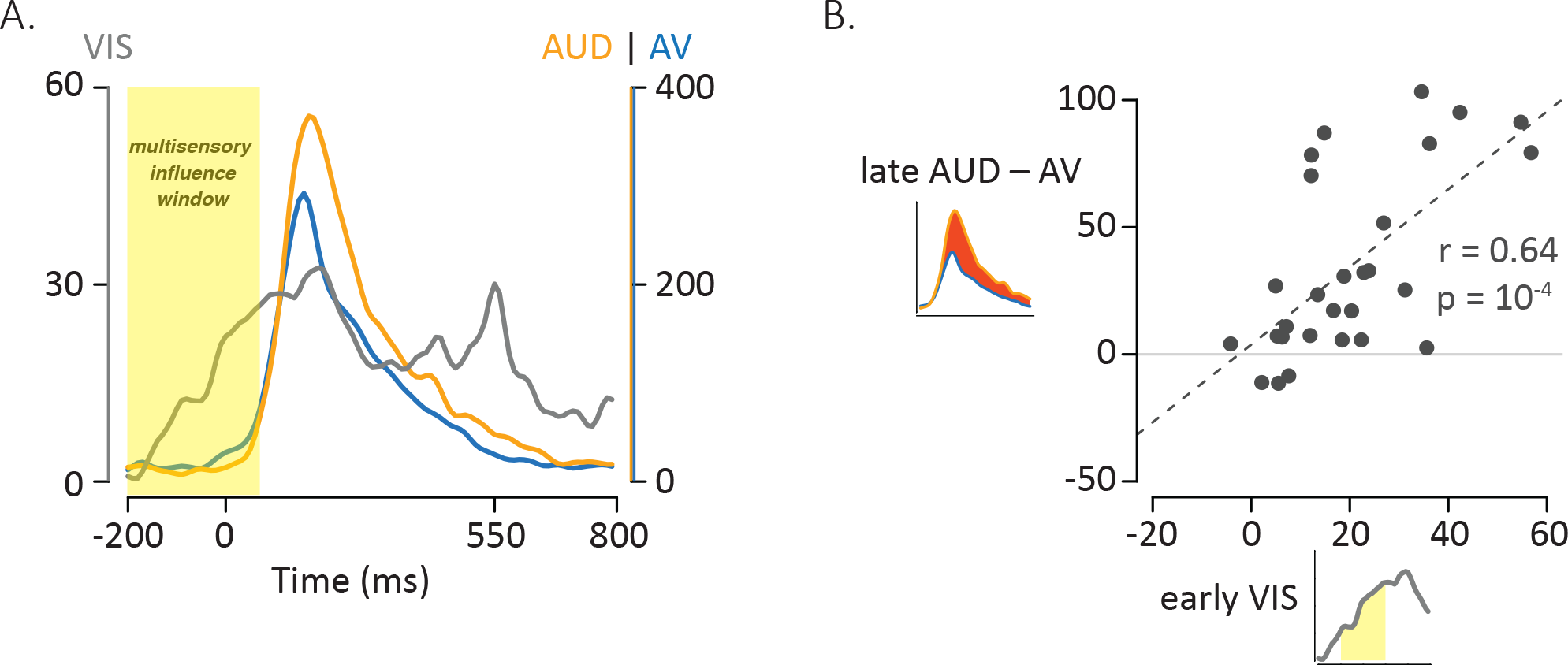
Influence of Visual Speech on Multisensory Integration. A. Responses to the three formats of mouth-leading words (visual-only, gray; auditory-only, orange; audiovisual, blue). Responses were aligned to auditory onset at *t* = 0 (or to the time when auditory onset would have occurred for visual-only stimuli). The left vertical axis contains the scale for visual-only neural responses (0% to 60%), which were much smaller than auditory-only and audiovisual responses (scale given in right-hand vertical axis; 0% to 400%). The visual-only response onset occurred prior to auditory-only response onset, creating a multisensory influence window (yellow region) in which visual speech could influence processing of auditory speech, resulting in a reduced response to audiovisual compared with auditory-only speech. B. The amplitude of the early neural response (BHA) to visual-only stimuli was positively correlated with the difference in the neural response between audiovisual and auditory-only speech (N = 28; *r* = 0.64, *p* = 10^−4^). The early visual-only response (horizontal axis) for each electrode was the average BHA for the 200-ms period following visual speech onset (−100 ms to 100 ms; yellow region in axis inset). The difference between the audiovisual and auditory-only neural response (vertical axis) was calculated as the difference in average BHA during auditory speech (0 ms to 550 ms; red region in axis inset).

In the conceptual model, visual speech information exerts a multisensory influence on auditory speech representations. Therefore, stronger responses to visual speech might result in more powerful multisensory interactions. To test this idea, we calculated the amplitude of the early neural response to visual-only words (0 to 200 ms following the onset of the first visual mouth movement) and compared it with the multisensory effect, defined as amplitude of the reduction for audiovisual *vs.* auditory-only words. As shown in Figure 4B, there was a significant positive relationship across electrodes between early visual response and multisensory influence (r = 0.64, *p* = 10^−4^).

### Multisensory Influence of Visual Speech: Voice-Leading Words

For voice-leading words, the conceptual model makes very different predictions. For these words, visual speech does not begin before auditory speech, leaving less opportunity for multisensory influence. To test this prediction, we examined the neural response to voice-leading words. There was again a robust neural response to visual-only voice-leading words (Figure S1A), but the latency of the visual-only response was 140 ms later for voice-leading than for mouth-leading words (*p* = 0.0006) reflecting the physical timing of mouth movements in the stimuli, which are later for voice-leading words than for mouth-leading words (Figure 1). This meant that within the voice-leading words, the latency of the visual-only and auditory-only neural responses were similar (mean latency 10 ms later for visual-only *vs.* auditory-only speech, *p* = 0.77). As predicted by the model, this corresponded to little multisensory influence, measured as a similar response amplitude for voice-leading words regardless of whether they were presente.D. in the audiovisual or auditory-only format (129% *vs.* 134%, *p* = 0.4). Across electrodes, there was not a significant correlation between visual-only response amplitude and multisensory influence, defined as the difference between the audiovisual *vs*. auditory-only neural responses (Figure S1B; *r* = 0.18, *p* = 0.35).

### Control Analysis: Latency

Our analysis depended on accurate time-locking between stimulus presentation and neural response recording. A photodiode placed on the monitor viewed by the participants was used to measure the actual onset time of visual stimulus presentation, while a splitter duplicated the auditory output from the computer to measure the actual onset time of auditory stimulus presentation. Both signals were recorded by the same amplifier used to record neural data, ensuring accurate synchronization. Latencies were similar for auditory-only mouth-leading *vs.* voice-leading words (128 ms *vs.* 134 ms, *p* = 0.54) demonstrating that the alignment of responses to the physical onset of speech was effective.

### Control Analysis: Anatomical Specialization

Previous studies have described anatomical specialization within pSTG (Hamilton et al., 2018; Ozker et al., 2017). Electrodes were color coded by the amplitude of the multisensory influence, calculated as the difference between auditory-only and audiovisual mouth-leading words. When viewed on the cortical surface, no consistent organization of multisensory influence was observed (Figure S2).

## Discussion

It has long been known that viewing the talker’s face enhances the intelligibility of auditory speech (Sumby and Pollack, 1954). This phenomenon can be explained by a simple conceptual model in which the early arrival of visual speech suppresses the representation of incompatible phonemes and enhances the representation of compatible phonemes. To test the model, we took advantage of the natural variability between mouth-leading words, in which visual speech precedes auditory speech, and voice-leading words, for which the converse is true (Chandrasekaran et al., 2009; Schwartz and Savariaux, 2014). As predicted by the model, there was a greater multisensory benefit for the comprehension of mouth-leading words than for the comprehension of voice-leading words.

Mirroring the perceptual effect, in neural recordings from the pSTG there was a difference between audiovisual and auditory-only words for mouth-leading but not voice-leading words. Decreased neural responses for audiovisual compared with auditory-only mouth-leading words are attributable to a multisensory interaction between the auditory speech and the visual speech. Surprisingly, when visual speech was present for mouth-leading words, the perceptual accuracy *increased* but the neural response *decreased*. To understand this observation, we constructed a neural model (Figure 5). The neural model incorporated the conceptual model principles of suppression/enhancement by visual speech and the experimental observation that the pSTG contains populations of neurons that are partially selective for specific phonemes, resulting in reduced but non-zero responses for non-preferred phonemes (Hamilton et al., 2018; Mesgarani et al., 2014).

**Figure 5.**
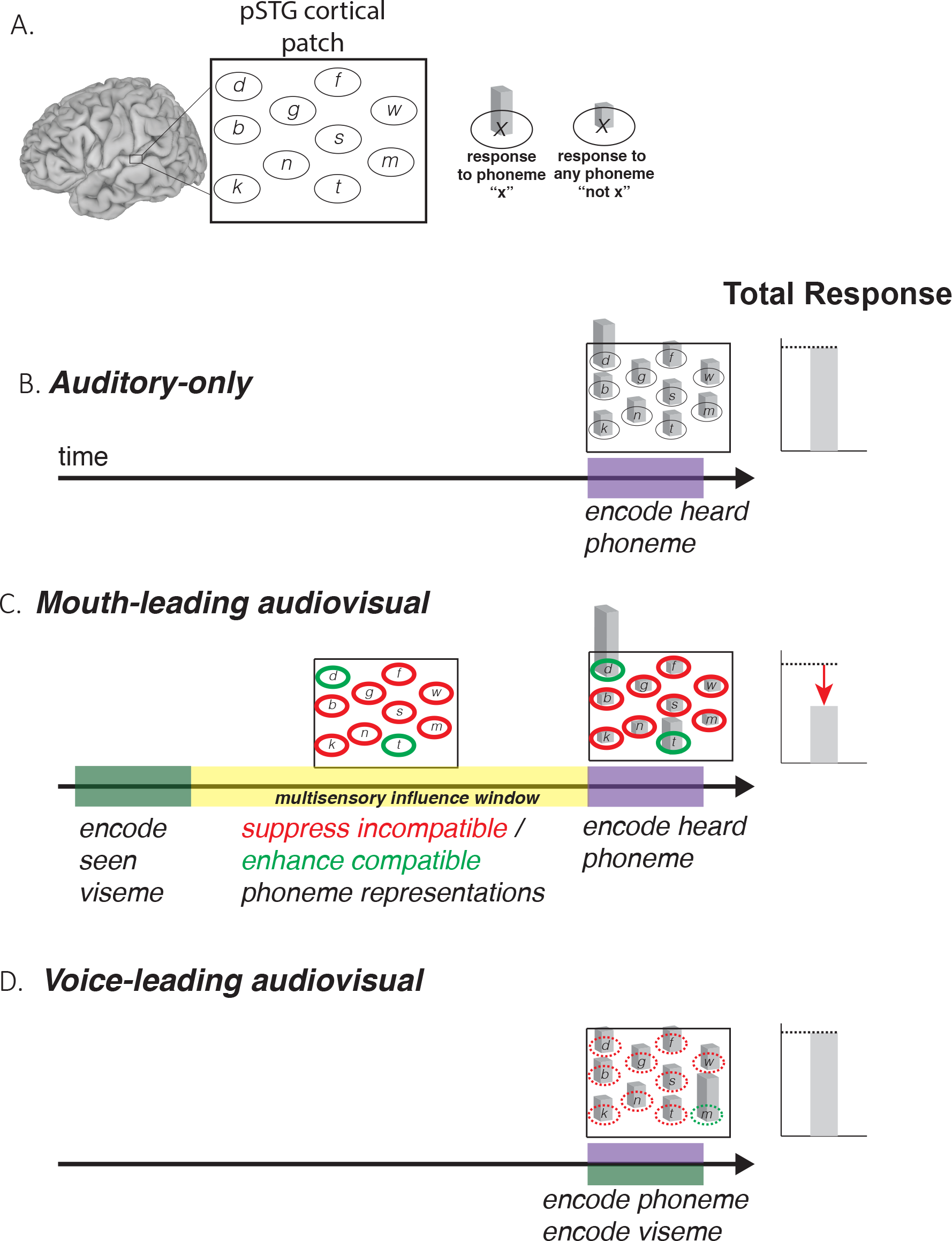
Model of Audiovisual Interactions in pSTG. A. In the pSTG, small populations of neurons are selective for specific speech sounds (phonemes). Each population is shown as an ellipse labeled by its preferred phoneme. The ellipses are shown spatially separated but the model is equally applicable if the neurons are intermixed instead of spatially segregated. Selectivity is only partial, so that for the population of neurons selective for a given phoneme “x” (ellipse containing “x”) presentation of the phoneme “x” evokes a large response (2 in our simple model), while presentation of any other phoneme (“not x”) evokes a small but non-zero response (1 in our model). B. When an auditory phoneme is presented, all populations of neurons respond, with the highest response in the population of neurons selective for that phoneme. Example shown is for presentation of auditory “d”; the amplitude of the response in each population of neurons is shown by the height of the bar inside each ellipse, with highest bar for “d” population. The total response summed across all populations is shown at right. C. For mouth-leading speech, early arriving visual speech suppresses activity in neuronal populations representing incompatible phonemes (red outlines) and enhances activity in neuronal populations representing compatible phonemes (green outlines). Arrival of auditory speech evokes activity in all populations, but activity is larger in compatible populations than incompatible populations. Example shown is for audiovisual “d”, resulting in larger responses in populations representing the compatible phonemes “d” and “t”, smaller responses in all other populations. The total response across all populations is decreased relative to the auditory-only format (dashed line and red arrow). D. For voice-leading speech, visual speech and auditory speech onset at similar times, resulting in no time for suppression or enhancement (dashed outlines; example shown is for audiovisual “m”). The total response is identical to the auditory-only format (dashed line).

Quantifying the neural model response required estimating the number of neuronal populations in pSTG, their selectivity, and the amplitude of suppression/enhancement. As a rough approximation, the model was constructed with 43 populations of neurons, one for each phoneme in the Jeffers and Barley classification scheme (Jeffers and Barley, 1971). Each population was assumed to respond with 2 arbitrary units of activity to the presentation of its preferred phoneme, and with 1 unit of activity to the presentation of any other phoneme. The amplitude of visual suppression was assumed to be ½ and the amplitude of visual enhancement was assumed to be two-fold. With these assumptions, presentation of the auditory-only syllable “d” resulted in 44 total units of activity (Figure 5B; 2 units in the population selective for /d/ and 1 unit in the other 42 populations) while presentation of the audiovisual syllable “d” resulted in 35.5 total units of activity, a decrease of 19% (Figure 5C; 4 units in the population selective for /d/, 2 units each in the 7 populations selective for other compatible phonemes, and 0.5 units each in the 35 populations representing incompatible phonemes). For voice-leading speech such as the audiovisual syllable “m” (Figure 5D) the model assumes that there is insufficient time for the suppression/enhancement of incompatible/compatible phonemes to manifest before the response evoked by the auditory phoneme, resulting in 44 total units of activity, identical to the auditory-only response.

As observed in the experimental data, the neural model predicted a *decreased* response to audiovisual compared with auditory-only speech for mouth-leading but not voice-leading words. The key factor in generating the decreased model response for mouth-leading words was the suppression of incompatible phoneme representations. Across a wide range of parameter values including the number of neuronal populations, the selectivity of each population, or the amount of enhancement/suppression, the model continued to predict reduced responses to audiovisual compared with auditory-only mouth-leading speech.

Both the conceptual and neural models assume enhancement of compatible populations as well as suppression of incompatible populations. However, only suppression was observed at the electrode level (in 25/28 pSTG electrodes; the remaining three electrodes showed similar responses to audiovisual and auditory-only speech). In all published classifications, there are more incompatible than compatible phonemes for any particular viseme (Cappelletta and Harte, 2012; Jeffers and Barley, 1971; Neti et al., 2000). This would lead to a predominance of suppression if pSTG electrodes recorded the total response summed across a combination of many suppressed populations and a few enhanced populations. Using smaller recording electrodes could allow for recording individual populations, separating those that show suppression from those that showed enhancement (Hamilton et al., 2018; Mesgarani et al., 2014). In the present study, we did not examine selectivity for different phonemes or visemes; both of these approaches will be important in future work.

### Relationship between neural suppression and perceptual enhancement

A property of the neural model is that visual speech does not activate phoneme representations directly but instead influences perception by modulating the activity evoked by auditory speech. This is consistent with perception. Silent viewing of the visual syllable “ga” does not produce an auditory percept, but when paired with the auditory syllable “ba”, many participant report hearing “da” (Basu Mallick et al., 2015; McGurk and MacDonald, 1976). Since the pSTG is known to be a brain hub for multisensory integration and audiovisual speech perception (Beauchamp, 2015) we make the parsimonious assumption that the perceptual and neural differences between mouth-leading and voice-leading words are related, although we did not directly compare them, as perception was measured with clear and noisy speech in healthy controls, while neural responses were measured only with clear speech in epileptic patients.

### Relationship to Predictive Coding

Predictive coding is a well-established principle at all levels of the auditory system (Denham and Winkler, 2017). Cross-modal suppression may result from similar mechanisms as predictive coding, with the difference that the information about the expected auditory stimulus does not come from previously-presented auditory stimuli but from early-arriving visual speech information. This expectancy is generated from humans’ developmental history of exposure to audiovisual speech, possibly through synaptic mechanisms such as spike-timing dependent plasticity (David et al., 2009). Over tens of thousands of pairings, circuits in the pSTG could be modified so that particular visual speech features inhibit or excite neuronal populations representing incompatible/compatible phoneme representations.

### Model Predictions and Summary

The conceptual model explains the absence of multisensory benefit for voice-leading speech because of the lack of a perceptual head start provided by visual speech, suggesting a number of interesting experiments. Voice-leading speech could be transformed by experimentally manipulating auditory-visual asynchrony, advancing the visual portion of the recording and rendering it effectively “mouth-leading” (Magnotti et al., 2013; Sánchez-García et al., 2018). Conversely, mouth-leading speech could be transformed by retarding the visual speech, rendering it effectively “voice-leading”. The model predicts that the transformed voice-leading speech would not exhibit neural cross-modal suppression and the concomitant perceptual benefit, while transformed mouth-leading speech would exhibit both features.

Our findings contribute to the grown literature of studies showing how visual input can influence the auditory cortex, especially pSTG (Besle et al., 2008; Ferraro et al., 2019; Kayser et al., 2008; Megevand et al., 2019; Zion Golumbic et al., 2013). Together with previous work showing that visual cortex is modulated by the presence or absence of auditory speech (Schepers et al., 2015), audiovisual speech is a prime example of how cross-modal interactions are harnessed by all levels of the cortical processing hierarchy in the service of perception and cognition (Ghazanfar and Schroeder, 2006).

## Methods

### Human Subject Statement

All experiments were approved by the Committee for the Protection of Human Subjects at Baylor College of Medicine.

### Stimuli

The stimuli in all experiments consisted of two exemplars of mouth-leading speech (“drive” and “last”) and two exemplars of voice-leading speech (“meant” and “known”). To estimate the auditory-visual asynchrony in the different stimuli, Adobe Premier was used to analyze individual video frames (30 Hz frame rate) and the auditory speech envelope (44.1 kHz). Visual onset was designated as the first video frame containing a visible mouth movement related to speech production. Auditory onset was designated as the first increase in the auditory envelope corresponding to the beginning of the speech sound. These values were: “drive” 170 ms/230 ms (visual onset/auditory onset); “last” 170ms/270ms; “meant” 170ms/130ms; “known” 200ms/100ms.

To visualize the complete time course of auditory and visual speech, shown in Figure 1, two of the stimuli were re-recorded at a video frame rate of 240 Hz (these re-recorded stimuli were not used experimentally). The instantaneous mouth size was determined in each video frame using custom software written in R (R Core Team, 2017) that allowed for manual identification of 4 control points defining the bounding box of the talker’s mouth. The area of this polygon was calculated in each frame and plotted over time. Auditory speech was quantified as the volume envelope over time, calculated by extracting the auditory portion of the recording, down-sampling to 240 Hz, and calculating the absolute value of the amplitude at each time step. The visual and auditory speech values were individually smoothed using a cubic spline curve with 30 degrees of freedom.

### Perceptual Data Analysis and Collection

The goal of the perceptual data analysis was to determine if the addition of visual information improved perception differently for mouth-leading and voice-leading words. Statistically, this was determined by testing the interaction (difference of differences) between word format (audiovisual *vs.* auditory-only) and word type (mouth-leading *vs.* voice-leading). While interactions can be tested with ANOVAs, accuracy data are proportional (bounded between 0% and 100%), violating the assumptions of the test. Instead, we applied a generalized linear mixed-effects model (GLMM) using odds-ratios (proportional change in accuracy, defined as the ratio of the probability of a correct response to the probability of an incorrect response) rather than absolute accuracy differences. For instance, an accuracy increase from 5% to 15% (absolute change of 10%) corresponds to a 3.4-fold increase in the odds-ratio (0.05/0.95 : 0.15/0.85) while an accuracy increase from 50% to 60% (absolute change of 10%) corresponds to a 1.5-fold odd-ratio increase (0.5/0.5 : 0.6/0.4).

Perceptual data and R code used for the data analysis and power calculations are available at https://openwetware.org/wiki/Beauchamp:DataSharing#Cross-modal_Suppression.

Power was calculated using parameters from a previous study with similar methods (Rennig et al., 2018). Each participant was assigned a randomly-selected accuracy level for understanding auditory noisy speech, ranging from 0% to 50% across participants; adding visual speech increased accuracy within each participant by 30% for mouth-leading words and by 20% for voice-leading words. We sampled a binomial distribution using the actual experimental design, with 40 participants and 20 trials for each of the four conditions. The critical test was for the interaction within the GLMM between word type (mouth-leading and voice-leading) and format (auditory-only and audiovisual). In 10,000 boot-strapped replications, the power to detect the simulated 10% interaction effect was 80%.

Perceptual data and R code used for the data analysis and power calculations are available at https://openwetware.org/wiki/Beauchamp:DataSharing#Cross-modal_Suppression.

46 participants were presented with the four stimulus exemplars using Amazon Mechanical Turk (https://www.mturk.com/). Within each participant, each exemplar was presented in four different formats: auditory-only (5 trials); auditory-only with added auditory noise (10 trials); audiovisual (5 trials); audiovisual with added auditory noise (10 trials). The trials were randomly ordered, except that in order to minimize carry over-effects all noisy stimuli were presented before any clear stimuli were presented. To construct the stimuli from the original audiovisual recordings, the auditory track of each exemplar was extracted using Matlab and all tracks were normalized to have similar mean amplitudes. To add auditory noise, white noise was generated with the same mean amplitude; the amplitude of the speech stimuli were reduced by 12 dB; the speech and noise auditory signals were combined; and the modified auditory track was paired with the visual track (for noisy audiovisual format). After each trial, participants responded to the prompt *“Type an answer in the text box to indicate what you heard. If you are not sure, take you best guess.”* No feedback was given. Six participants had very low accuracy rates (from 0% to 75%) for clear words, suggesting that they were not attending to the stimuli. These participants were discarded, resulting in an *n* of 40 (adequate given the power calculation described above).

Because participant responses were collected using a text-box (free-response) preprocessing was required before analysis. Spelling and typographical errors were corrected. Responses were classified as correct based on whether the initial viseme of the typed response matched the initial viseme of the presented word using the Jeffers classification system (Jeffers and Barley, 1971). Analysis was conducted in R (R Core Team, 2017) using the *glmer* function (family set to binomial) from the *lme4* package (Bates et al., 2015). The initial GLMM contained fixed effects of word format (auditory-only *vs.* audiovisual) and word type (mouth-leading *vs.* voice-leading), the word format-by-word type interaction, and a random effect for participant (different intercept for each participant) and exemplar. The baseline for the model was set to the response to mouth-leading words in the auditory-only word format. The inclusion of random effects allowed for participant differences but meant that the estimated odds-ratios are different than those calculated from the raw accuracy score.

For further analysis, separate GLMMs were created for each word type, with a fixed effect of word format (auditory-only *vs.* audiovisual), random effect of participant and baseline set to auditory-only word format. These separate GLMMs were used to calculate the reported odds-ratio and significance within word type.

### Neural Studies

Eight subjects (5F, mean age 36, 6L hemisphere) who were selected to undergo intracranial electrode grid placement for phase 2 epilepsy monitoring provided informed consent to participate in this research protocol. Electrode grids and strips were placed based on clinical criteria for epilepsy localization and resection guidance. The research protocol was approved by the Baylor College of Medicine Institutional Review Board. After a short recovery period following electrode implantation, patients were presented with audiovisual stimuli while resting in their hospital bed in the epilepsy monitoring unit. Stimuli were presented with an LCD monitor (Viewsonic VP150, 1024 x 768 pixels) placed 57cm in front of the subject’s face, and sound was projected through two speakers mounted on the wall behind and above the patient’s head.

The four stimulus exemplars were presented in three formats: auditory-only, visual-only, and audiovisual. Auditory-only trials were created by replacing the speaker’s face with a blank screen consisting only of a fixation target. Visual-only trials were created by removing the auditory component of the videos. No auditory noise was present in any format. Stimuli were presented in random interval. The behavioral task used a catch trial design. Subjects were instructed to respond only to an audiovisual video of the talker saying “press”. No feedback was given. Neural data from catch trials was not analyzed.

### Neurophysiology Recording and Data Preprocessing

Implanted electrodes consisted of platinum alloy discs embedded in flexible silastic sheets (Ad-Tech Corporation, Racine, WI). Electrodes with both 2.3 mm and 0.5 mm diameter exposed surfaces were implanted, but only electrodes with 2.3 mm were included in this analysis. After surgery, electrode tails were connected to a 128-channel Cerebus data acquisition system (Blackrock Microsystems, Salt Lake City, UT) and recorded during task performance. A reversed intracranial electrode facing the skull was used as a reference for recording, and all signals were amplified, filtered (high-pass 0.3 Hz first-order Butterworth, low pass 500 Hz fourth-order Butterworth), digitized at 2000 Hz, then converted from Blackrock format to MATLAB (MathWorks Inc. Natick, MA). Stimulus videos were presented using Psychtoolbox software package (Brainard 1997; Pelli 1997; Kleiner, Brainard, and Pelli 2007) for MATLAB. The auditory signal from the stimulus videos was recorded on a separate channel simultaneously and synchronously along with the electrocorticography voltage.

Preprocessing was performed using the R Analysis and Visualization of intracranial Electroencephalography (RAVE) package (openwetware.org/wiki/Beauchamp:RAVE). Data was notch filtered (60 Hz and harmonics), referenced to the average of all valid channels, and converted into frequency and phase domains using a wavelet transform. The number of cycles of the wavelet was increased as a function of frequency, from 3 cycles at 2 Hz to 20 cycles at 200 Hz, to optimize tradeoff between temporal and frequency precision (Cohen, 2014). Data was down-sampled to 100 Hz after the wavelet transform. The continuous data was epoched into trials using the auditory speech onset of each stimulus as the reference (*t* = 0). For visual-only trials, *t* = 0 was considered to be the same time at which the auditory speech would have begun, as determined from the corresponding audiovisual stimulus.

### Calculation of Broadband High-Frequency Activity (BHA)

For each trial and frequency, the power data were transformed into percentage signal change from baseline, where baseline was set to the average power of the response from −1.4 to −0.4 seconds prior to auditory speech onset. This time window consisted of the inter-trial interval, during which participants were shown a gray screen with a white fixation point. The percent signal change from this pre-stimulus baseline was then averaged over frequencies from 75-150 Hz to calculate the broadband high-frequency activity (BHA). Trials with median absolute differences more than five standard deviations from the mean were excluded (<2% excluded trials). For visualization in Figure 3A, the average BHA for auditory-only trials during auditory speech (time window 0.0 to 0.55 seconds) was calculated for each electrode and compared against baseline (single sample *t*-test). The resulting *p*-value was plotted according to a white to orange color scale (white is *p* = 0.001, Bonferroni-corrected; orange is *p* < 10^−14^)

### Electrode Localization and Selection

FreeSurfer (Dale et al., 1999; Fischl et al., 1999) was used to construct cortical surface models for each subject from their preoperative structural T1 magnetic resonance image scans. Post-implantation CT brain scans, showing the location of the intracranial electrodes, were then aligned to the preoperative structural MRI brain using the Analysis of Functional Neuroimaging (AFNI; Cox, 1996) package and electrode positions were marked manually on the structural surface model. Electrode locations were projected to the surface of the cortical model using AFNI. SUMA (Argall et al., 2006) was used to visualize cortical surface models with overlaid electrodes, and positions were confirmed using intraoperative photographs of the electrode grids overlaid on the brain when available. Cortical surface models with overlaid electrodes were mapped to the Colin N27 brain (Holmes et al., 1998) using AFNI/SUMA, allowing for visualization of electrodes from all subjects overlaid over one single brain atlas.

All analyses were performed on electrodes (*n* = 28 from eight participants; Figure 3A) that met both an anatomical criterion (located over the posterior superior temporal gyrus) and a functional criterion (significant BHA response to auditory-only speech). The anatomical border between anterior and posterior superior temporal gyrus was defined by extending a line inferiorly from the central sulcus to split the superior temporal gyrus into anterior and posterior portions. The functional criterion was a significant (*p* < 0.001, Bonferroni-corrected) BHA response to the auditory-only word format during the period from stimulus onset to stimulus offset (0 seconds to 0.55 seconds). Because the functional criterion ignored word type (voice-leading *vs.* mouth-leading) and did not include audiovisual stimuli, the main comparison of interest (the interaction between response amplitude for different word types and stimulus word formats) was independent of the functional criterion and hence unbiased.

### Statistical Analysis of Neural Data

Neural responses were collapsed into a single value by averaging the high-frequency activity (BHA) across the time window from stimulus onset to stimulus offset (0-s to 0.55 seconds) for each trial for each electrode. These values were then analyzed using a linear mixed-effects model (LME) with a baseline of mouth-leading words in the auditory-only word format. The model factors were two fixed effects of word format (auditory-only *vs.* audiovisual) and word type (mouth-leading *vs.* voice-leading), a word format-by-word type interaction was included, as well as random effects of electrode nested within subject (random intercepts and slopes for each subject-by-electrode pair). Because electrodes were selected based on the response to auditory-only speech ignoring word type (regardless of response to audiovisual or visual-only speech), the model was unbiased. All LMEs were performed in R using the *lmer* function in the *lme4* package. Estimated *p*-values were calculated using the Satterthwaite approximation provided by the *lmerTest* package (Kuznetsova et al., 2017).

To further explore the word format-by-word type interaction, we created separate LMEs for each word type (mouth-leading and voice-leading). For each word type, the LME had fixed effect of word format (auditory-only *vs.* audiovisual) and random effects of electrode nested within subject, with baseline set to the auditory-only word format. These separate LMEs were used to calculate the significance and magnitude of the effect of auditory-only and audiovisual word format on BHA in pSTG within word type.

For the analysis shown in Figure 4, the average BHA for each visual-only word format trial was calculated over the time window from −0.1 to 0.1 seconds, the time interval when visual speech was present for mouth-leading words but absent for voice-leading words. We then created an LME model with fixed effect of word type and random effects of electrode nested within subject. The neural response onset stim was measured by calculating the average time (across trials) it took the BHA to reach half its maximum value for each word format and word type. Paired *t*-tests were used to compare half-maximum times between specific word types and word formats.

## Acknowledgments

This research was supported by NIH R01NS065395 and R25NS070694.

---

**Figure S1.**
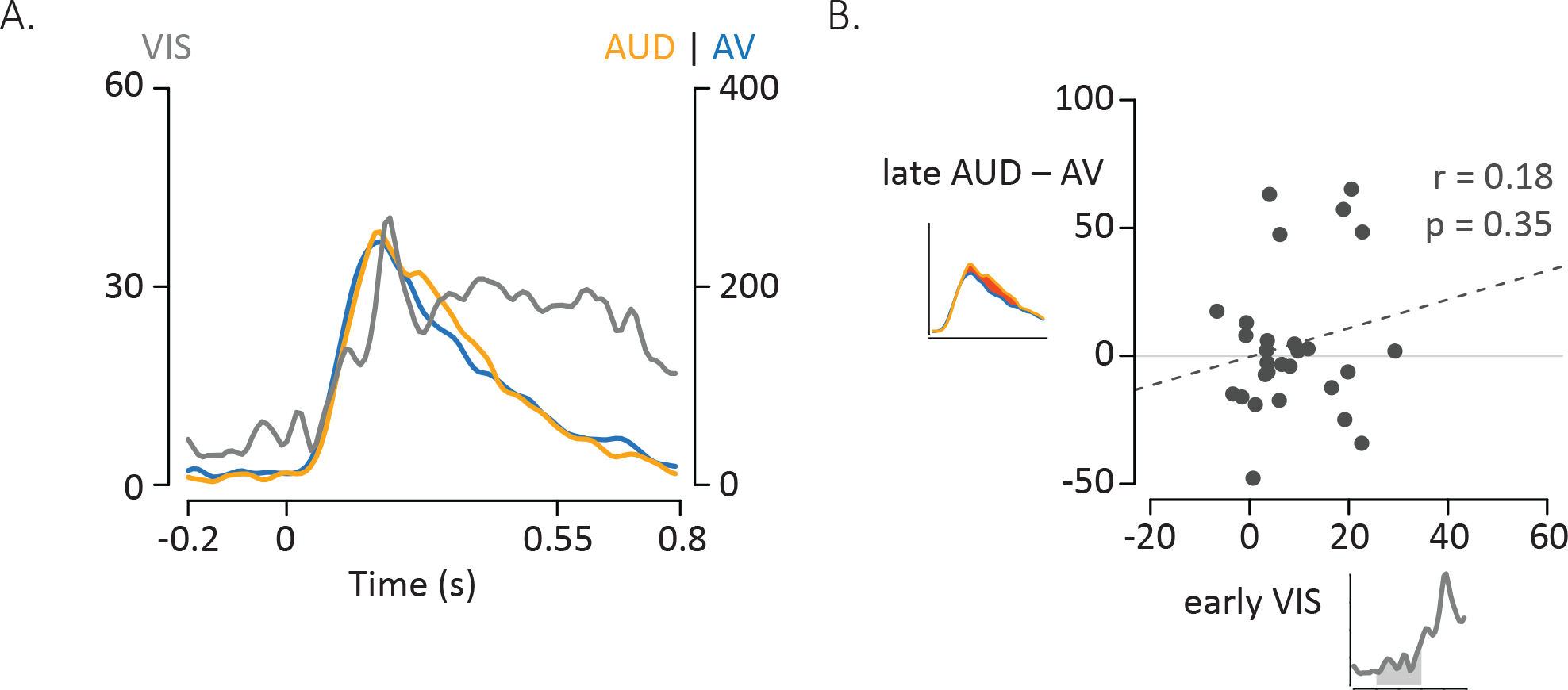
Influence of Visual Speech for Voice-Leading Words. A. Responses to the three formats of voice-leading words (visual-only, gray; auditory-only, orange; audiovisual, blue). Responses were aligned to auditory onset at *t* = 0 (or to the time when auditory onset would have occurred for visual-only stimuli). The left vertical axis contains the scale for visual-only neural responses (0% to 60%), which were much smaller than auditory-only and audiovisual responses (scale given in right-hand vertical axis; 0% to 400%). The neural response to the three word began at similar times. B. Correlation between the amplitude of the early neural response to visual-only words and the difference in the neural response between audiovisual and auditory-only speech. The early visual-only response (horizontal axis) for each electrode was the average BHA for the 200-ms period following visual speech onset (time −100 ms to 100 ms; grey region underneath curve in axis inset). The reduction in neural response to audiovisual *vs.* auditory-only speech (vertical axis) was calculated as the difference in average BHA during the duration of the entire auditory stimulus (0 ms to 550 ms; red region in axis inset).

**Figure S2.**
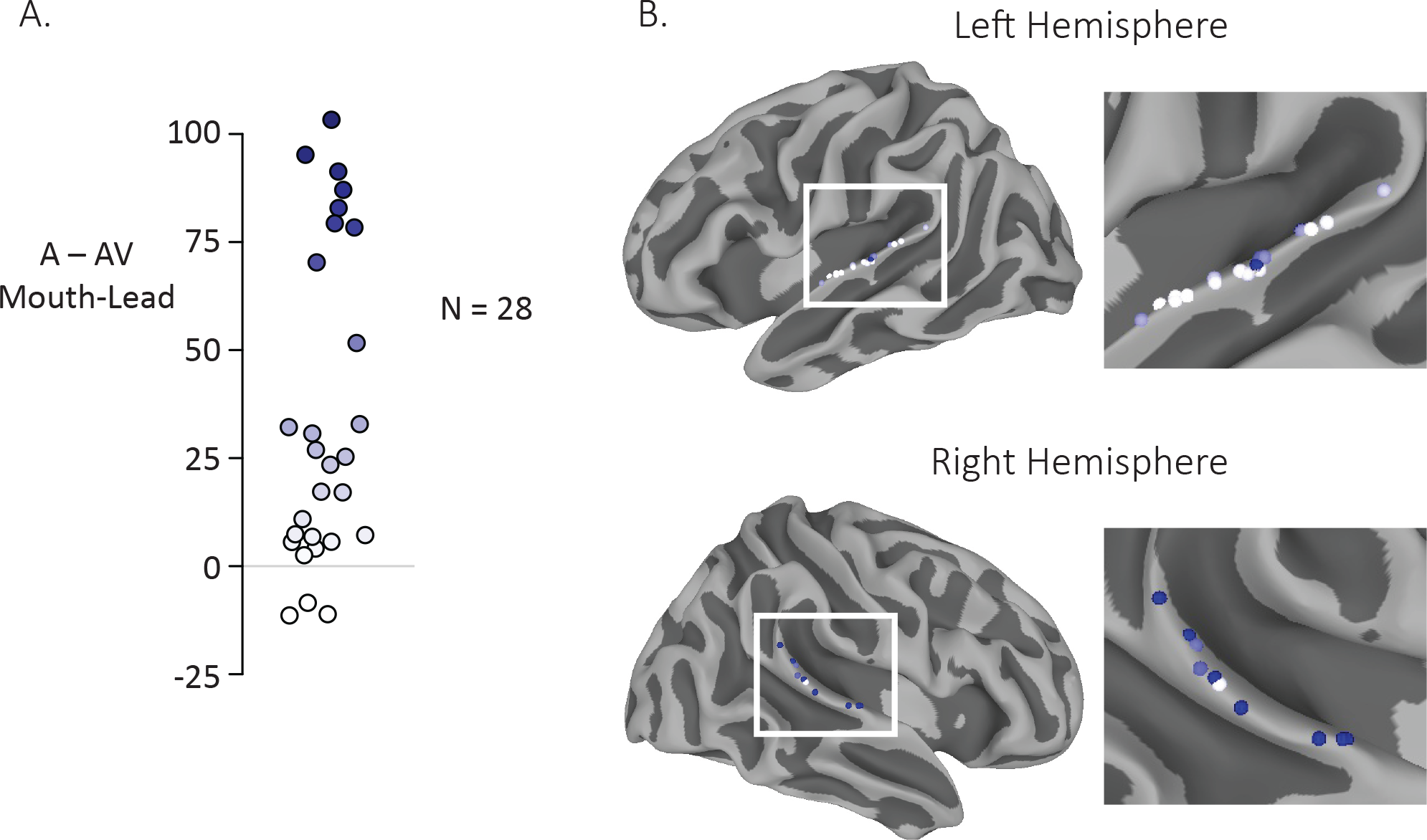
Anatomic distribution of multisensory gain by electrode. A. The reduction in neural response to audiovisual compared to auditory-only speech for mouth-leading words was measured for each electrode (difference in average BHA to audiovisual stimuli and average BHA to auditory-only stimuli over time window 0 ms to 550 ms). Most electrodes (25 of 28) had a decreased neural response to audiovisual compared to auditory only stimuli. Color on the white-to-blue gradient corresponds to amount of reduction. B. All electrodes from all participants displayed on a template brain (same color scale as A). No consistent organization of multisensory influence was observed.

